# Medial temporal lobe structure, mnemonic and perceptual discrimination in healthy older adults and those at risk for mild cognitive impairment

**DOI:** 10.1101/2022.05.13.491810

**Authors:** Helena M. Gellersen, Alexandra N. Trelle, Benjamin G. Farrar, Gillian Coughlan, Saana M. Korkki, Richard N. Henson, Jon S. Simons

## Abstract

Cognitive tests sensitive to the integrity of the medial temporal lobe (MTL), such as mnemonic discrimination of perceptually similar stimuli, may be useful early markers of risk for cognitive decline in older populations. Perceptual discrimination of stimuli with overlapping features also relies on MTL, but remains relatively unexplored in this context. We assessed mnemonic discrimination in two test formats (Forced Choice, Yes/No) and perceptual discrimination of objects and scenes in 111 community-dwelling older adults at different risk status for cognitive impairment based on neuropsychological screening. We also investigated associations between performance and MTL subregion volume and thickness. The at-risk group exhibited reduced entorhinal thickness and impaired perceptual and mnemonic discrimination. Perceptual discrimination impairment partially explained group differences in mnemonic discrimination and correlated with entorhinal thickness. Executive dysfunction accounted for Yes/No deficits in at-risk adults, demonstrating the importance of test format for the interpretation of memory decline. These results suggest that perceptual discrimination tasks may be useful tools for detecting incipient cognitive impairment related to reduced MTL integrity in non-clinical populations.

## Introduction

The ability to form high-fidelity representations of visual stimuli is reliant on the medial temporal lobe (MTL) and is essential to distinguish similar inputs during both perception and memory (Barense et al., 2007; Bussey & Saksida, 2002; Graham et al., 2010). The MTL is vulnerable to normal age-related structural and functional change (Fjell et al., 2014; Leal & Yassa, 2013). This is thought to arise, in part, due to the accumulation of neurofibrillary tangle pathology beginning in the entorhinal cortex (ERC) that is a hallmark feature of Alzheimer’s disease (Braak & Braak, 1995). As a result, with increasing age and during the course of Alzheimer’s disease (AD), perceptual and mnemonic representations become less detailed to the detriment of a variety of cognitive processes (Koen et al., 2019; Koen & Rugg, 2019; Leal & Yassa, 2014; A. C. H. Lee et al., 2007; Trelle et al., 2019). Tests that require discrimination between stimuli with high degrees of feature overlap, such as the mnemonic similarity task (Stark et al., 2019), may be useful tools to probe the integrity of perirhinal-entorhinal-hippocampal circuits and hold promise for the identification of community-dwelling older adults who may be at increased risk of AD as well as for tracking progressive MTL degeneration and evaluating efficacy of new disease-modifying treatments and interventions (Adams et al., 2020; Ally et al., 2013; Berron et al., 2019; Gellersen, Coughlan, et al., 2021; Holden et al., 2013; Leal et al., 2019; S. Lee et al., 2020; Sinha et al., 2018; Trelle et al., 2021; Webb et al., 2020).

Because the MTL regions vulnerable to AD pathology are also involved in complex perception (Graham et al., 2010), it is possible that individuals at risk for MCI may not only be impaired on mnemonic discrimination of highly similar stimuli, but may also show deficits in perceptual discrimination when tasks require distinguishing between similar complex stimuli (Yeung et al., 2017, 2019). Due to similar demands on MTL representations, individuals with MCI risk may be impaired in both tasks, yet, to date perceptual discrimination has been largely unexplored in this context. Here we explore how task demands might modulate differences in perceptual and mnemonic discrimination across risk groups. Specifically, our first aim explores the role of test format (Yes/No, Forced Choice) on mnemonic discrimination performance and stimulus category (object, scene) and feature ambiguity (high, low) on perceptual discrimination performance. Our second aim explores perceptual discrimination and executive function as potential mediators of mnemonic discrimination performance across test formats. Our third aim explores differences in MTL subregional integrity across groups, as well as associations between subregional integrity and perceptual-mnemonic discrimination performance.

The question of how task demands influence mnemonic discrimination impairments in older adults at risk for MCI is not well understood. One dimension that is likely to be important is the level of demand for strategic retrieval processes, that differ, for example, in performance on Forced Choice versus Yes/No task formats of recognition memory tests with similar lures (Gellersen, Trelle, et al., 2021). To date, most studies have used variants of the Yes/No format, such as Old/Similar/New judgments (Stark et al., 2019), in which one item is presented at a time, and participants need to impose a criterion of evidence in order to choose one response category. These studies reported that cognitively normal older adults with biomarker evidence for AD pathology were significantly impaired (Berron et al., 2019; Trelle et al., 2021). However, this Yes/No task format places substantial demands on strategic retrieval, such as the selection of response criteria and use of “recall-to-reject” strategies for recollection to reduce false alarms (Cohn et al., 2008; Gallo, 2004; Gallo et al., 2006; Migo et al., 2009; Trelle et al., 2017). As a result, executive dysfunction, which is also common in ageing and related to atrophy in prefrontal cortex (PFC), may be a significant driver of performance in tests using this format (Gellersen, Trelle, et al., 2021). In contrast, the Forced Choice format provides retrieval support by presenting both a target and a lure simultaneously on each test trial. Tests of this type can be solved by comparing the relative levels of evidence for each choice, without imposing an absolute response criterion, and therefore have lower demands on executive functioning. Moreover, it has been suggested that Forced Choice tasks can be performed on the basis of differences in memory strength or relative familiarity between exemplars, rather than recollection of specific details, even when lures are perceptually similar (Angel et al., 2013; Gellersen et al., 2021; Holdstock et al., 2002; Migo et al., 2009; Migo et al., 2014; Trelle et al., 2017), therefore reducing demands on hippocampal and frontal processes in support of recollection-based retrieval and increasing the relative contribution of perirhinal (PRC) and entorhinal cortices (ERC) (Bowles et al., 2007; Holdstock et al., 2002; Vann et al., 2009). By contrasting performance in Forced Choice and Yes/No tests, we test whether the association between MCI risk and mnemonic discrimination is dependent on the degree to which a task taxes hippocampal memory processes.

We view mnemonic discrimination deficits in older adults through the lens of a representational account, in which target-lure similarity is one key determinant of impairment, consistent with the representational-hierarchical framework of MTL function (Bussey & Saksida, 2002; Cowell et al., 2010; Kent et al., 2016). This framework proposes that formation of complex, conjunctive visual representations necessary to differentiate stimuli with overlapping features is dependent on MTL, and lesions to as well as normal age-related and early pathological changes in these regions result in impoverished visual representations that compromise any cognitive process reliant on them, whether that be mnemonic or perceptual (Barense et al., 2005; Burke et al., 2018; A. C. H. Lee et al., 2007). If the quality of these representations were impaired, mnemonic discrimination deficits among individuals with early signs of cognitive decline would therefore be predicted in both Yes/No and Forced Choice tests. In line with the proposal that representational quality may underpin mnemonic discrimination abilities, age-related declines in mnemonic discrimination have been linked to performance on perceptual tasks that require discrimination of simultaneously-presented, highly-similar objects or scenes in tasks without long-term memory demands (Gellersen, Trelle, et al., 2021; Trelle et al., 2017). These deficits are particularly prominent under conditions of high feature overlap, perceptual interference and demands on viewpoint-invariant processing which crucially relies on MTL (Burke et al., 2012; Gellersen, Trelle, et al., 2021; Newsome et al., 2012). The representational-hierarchical view of MTL function offers one explanation as to why age-related deficits are often present in Forced Choice recognition memory tests under conditions of high target-lure similarity but may be absent when distinguishing between targets and novel foils: reduced availability of item details will be detrimental to mnemonic discrimination regardless of demands on strategic retrieval (Bastin & van der Linden, 2003; Gellersen, Trelle, et al., 2021; Koen & Yonelinas, 2014; Trelle et al., 2017; Yonelinas, 2002). Following from this representational-hierarchical view, a loss of integrity of ERC and PRC-dependent functions will ultimately be detrimental to performance on any mnemonic discrimination task. As a result we expect that older adults at risk for MCI will be impaired across both Forced Choice and Yes/No tasks and that a proxy for complex perceptual processing will be a predictor of mnemonic function in this sample (Gellersen, Trelle, et al., 2021; Trelle et al., 2017). Of note, in our analyses on the association between perceptual processes and mnemonic discrimination, we refer to these higher perceptual processes rather than basic properties of the visual system such as acuity and contrast sensitivity which have also previously been shown as important for mnemonic discrimination (Davidson et al., 2019).

Moreover, we expect that these deficits in the discrimination of highly similar visual stimuli would not be restricted to the memory domain but extend to perceptual processes as well. Few studies have explored perceptual discrimination of stimuli with overlapping features in populations at risk for cognitive impairment. However, prior work has demonstrated that perceptual discrimination of highly similar abstract objects is impaired under conditions of high interference in MCI and in individuals who are at-risk for MCI (Newsome et al., 2012) as well as in middle-aged individuals at genetic risk for AD (Mason et al., 2017). This decline in perceptual discrimination ability may be capable of (at least partially) accounting for mnemonic discrimination deficits in older adults at risk for MCI. Indeed, in a sample of non-clinical older adults that included individuals who failed the Montreal Cognitive Assessment (a neuropsychological screening tool to detect MCI) (Nasreddine et al., 2005), lower anterolateral entorhinal cortex volume was associated with a decreased preference to view similar but novel as opposed to repeated stimulus configurations (Yeung et al., 2017), suggesting altered processing of complex perceptual conjunctions as a function of MCI risk. Taken together, we expect older adults with early signs of cognitive decline to be impaired on all tasks that share common demands on the MTL, regardless of whether they are needed to support perceptual or mnemonic processes (Braak & Braak, 1991; Dickerson & Sperling, 2009; Holbrook et al., 2019; Olsen et al., 2017; Speer & Soldan, 2015; Wolk et al., 2013).

Our third aim leverages MRI measures of MTL subregional grey matter integrity, including both volumes of MTL subregions derived from manual segmentations of high-resolution images and cortical thickness estimated from automated segmentation. We investigate whether MTL integrity differs across groups, is correlated with individual differences in perceptual and mnemonic discrimination, and whether brain-behaviour relationships vary as a function of task demands and stimulus category. Prior studies have assessed the relationship between mnemonic discrimination and MTL subregions and found associations with CA3 and dentate gyrus volume (Bennett et al., 2019; Stark & Stark, 2017). To our knowledge, no studies have explored how this association might vary as a function of task demands, nor how MTL subregion volume relates to complex perceptual discrimination in older adults without dementia diagnosis.

This work is predicated on research suggesting that the various MTL subregions have distinct roles in specific memory processes and in the processing of different visual stimulus categories (Argyropoulos et al., 2021; Graham et al., 2010; Reagh et al., 2014; Reagh & Yassa, 2014). We include both volumes of MTL subregions derived from manual segmentations of high-resolution images and cortical thickness estimated from automated segmentation. These analyses allow us to test whether tasks known to be sensitive to MTL subregional integrity are also sensitive to individual variability in MTL subregional volume in older adults without clinical memory impairment. We formulate the following predictions regarding brain-behaviour relationships between MTL grey matter and perceptual-mnemonic discrimination.

First, based on both functional and structural neuroimaging (Bennett et al., 2019; Stark & Stark, 2017; Yassa et al., 2011; Yassa & Stark, 2011), we predict that Yes/No performance will be associated with volume in hippocampal cornu ammonis 3 (CA3) and dentate gyrus (DG), which are essential for mnemonic discrimination in Yes/No and Old/Similar/New task formats due to their involvement in pattern separation. In contrast, we expect structural integrity of PRC to be a predictor of Forced Choice performance due to its importance in object-level representations and its role in familiarity-based memory judgments (Burke et al., 2018; Westerberg et al., 2013). While failure of pattern separation should be highly detrimental to Yes/No performance, a high-quality representation of the target item as formed by PRC should be able to support performance in the Forced Choice task, as demonstrated in prior work with hippocampal lesion subjects (Holdstock et al., 2002; Migo et al., 2009).

Second, it is well established that MTL lesions result in performance decrements in complex perceptual discrimination in a subregion-specific manner (Barense et al., 2005; Graham et al., 2006; A. C. H. Lee et al., 2007). We therefore may find differential contributions of MTL subregions to high ambiguity object and scene discrimination, respectively, with PRC volume being related to the former, and hippocampal and parahippocampal structural integrity to the latter. Finally, for ERC, we hypothesise that volumes and cortical thickness correlate with perceptual discrimination across stimulus categories and mnemonic discrimination scores across task formats (Yeung et al., 2017, 2019, 2021). We base this prediction on prior findings suggesting that the anterolateral and posterior-medial entorhinal subregions have been implicated in object and scene processing, respectively and that ERC is crucially involved in high-fidelity object perception and memory reinstatement (Berron et al., 2018; Charles et al., 2004; Schultz et al., 2012, 2019; Staresina et al., 2019). Given that we do not separate the ERC into its subregions, volume-behaviour associations may therefore be apparent regardless of stimulus material.

In summary, the present study investigates how risk for MCI impacts performance on mnemonic and perceptual discrimination tasks, including how performance is impacted by test format, stimulus category, and feature ambiguity and possible associations with MTL subregion structure. This approach allows us to assess whether older adults at risk for MCI are impaired across memory and perception tasks with similar demands on MTL representations and whether brain-behaviour relationships depend on task demands and stimulus category. Moreover, our analyses allow us to identify whether different cognitive functions such as complex perception and executive processes are potential mediators of the relationship between MCI risk and impairment in mnemonic discrimination as a function of task demands.

## Methods

### Participants

111 community-dwelling older adults (87 female) aged 60 to 87 (*M*=71.11, *SD*=5.40) participated in this study. Participants were native English speakers, had normal or corrected to normal vision, and no history of diagnosed psychiatric or neurological conditions. The study was approved by the Cambridge Psychology Research Ethics Committee.

Older adults were screened for cognitive impairment with the Montreal Cognitive Assessment (Nasreddine et al., 2005), an instrument used to measure global cognition. Of the 111 in the sample, 86 older adults performed within the normal range (score >= 26), whereas 25 older adults scored below the suggested cut-off (score <26; Damian et al, 2011). Here we classify the 25 older adults who performed below the normal range as at-risk for cognitive impairment, as in previous work using the MoCA (D’Angelo et al., 2016; Newsome et al., 2012; Olsen et al., 2017; Yeung et al., 2017, 2019). We refer to this group throughout the manuscript as the ‘at-risk group’ (AR). The group performing within the normal range is referred to as the ‘cognitively unimpaired group’ (CU). The at-risk group was significantly older than CU older adults (*t*(37.05)=3.09, *p*=.004). We therefore control for age in all analyses.

The 86 participants who passed the MoCA were also included in the analysis presented in Gellersen et al. (2021). Demographic and cognitive test data for both groups is summarised in Table 1.

**Table 1.** Demographics of the study sample stratified by MoCA score (at-risk vs. unimpaired).

| | Cognitively<br>unimpaired<br>(N=86) | At risk for MCI<br>(N=25) | $\beta$ |
| --- | --- | --- | --- |
| Sex | 49 female (57%) | 12 female (48%) |  |
| Age | 70.26 (5.11) | 74.04 (5.47) | -.70** |
| Education | 17.39 (4.95) | 16.50 (4.05) | .19 |
| MoCA Total (/30) | 28.08 (1.29) | 23.56 (1.45) | 4.46*** |
| MoCA Attention (/6) | 5.87 (.37) | 5.25 (.97) | .99*** |
| MoCA Visuospatial/Executive (/5) | 4.56 (.63) | 3.96 (.84) | .86*** |
| MoCA Language (/3) | 2.62 (.60) | 2.04 (.79) | .91*** |
| MoCA Memory (/5) | 4.10 (.99) | 1.84 (1.46) | 1.50*** |
| NART | 42.30 (4.80) | 37.52 (7.08) | .85*** |
| Shipley Vocabulary | 37.63 (1.32) | 35.84 (3.44) | .84*** |
| Trails A | 49.42 (17.33) | 59.92 (19.99) | -.38 |
| Trails B | 84.26 (25.38) | 116.40 (54.09) | -.76*** |
| Digit Span Total | 18.97 (3.97) | 15.36 (3.37) | .84*** |
| Verbal Fluency (Letters) | 16.12 (3.64) | 14.17 (4.82) | .46 |

Means ± Standard deviations. *F*: female. *P*-values for MoCA group effects are controlled for age. For one participant in the cognitively unimpaired group for whom years of education was not available, the missing entry was replaced by the mean educational attainment of the low-risk group. \* $p < .05$ , \*\* $p < .01$ , \*\*\* $p < .001$ .

### Behavioural tasks

Figure 1 presents a schematic of test stimuli and procedure (Gellersen, Trelle, et al., 2021; Trelle et al., 2017). The mnemonic discrimination task included 200 highly similar exemplar pairs with a target and a corresponding lure each. Good performance on this task relied on high fidelity representations of target objects. Participants viewed 200 items during study and to orient attention to the stimuli they were asked to indicate for each object whether it was bigger or smaller than a shoe box. Following a distractor task in which participants counted backwards from 100 in steps of seven, they were tested on their memory for objects in two separate test phases: 100 trials using a Forced Choice test format in which both target and lure were presented side by side, and 100 trials in the Yes/No task in which either the target or the lure were shown. Participants were instructed to select the old items and reject the similar lures. The order in which participants completed the two test formats was counterbalanced across participants.

**Figure 1.**
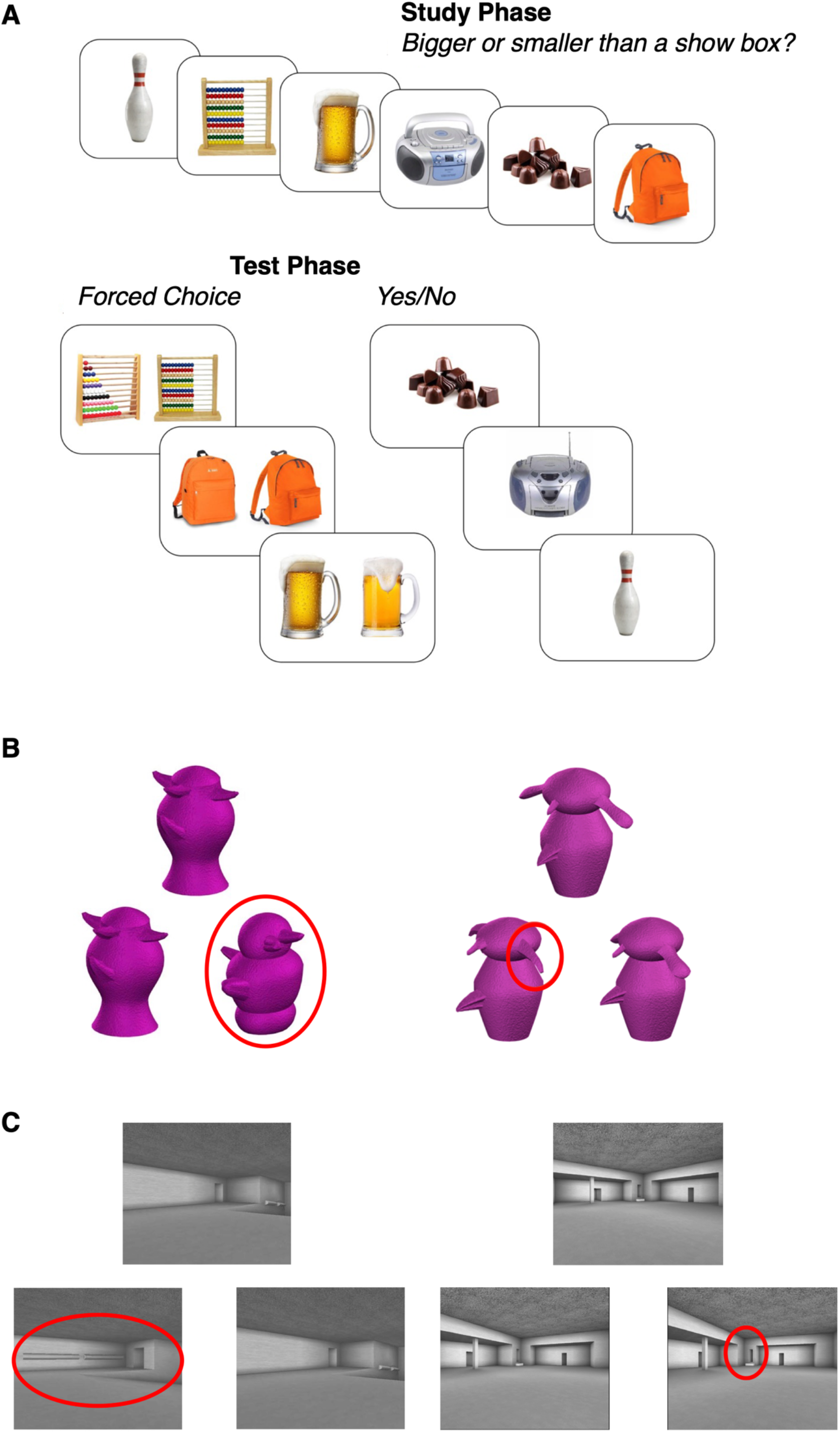
Schematic of the experimental paradigm. A. Mnemonic discrimination task. During study, participants were shown objects and made a size judgment for each object (top). In the test phase (bottom), previously studied objects were either shown next to new perceptually similar foil objects (Forced Choice task), or either the target or the foil object was shown (Yes/No task). B. Perceptual discrimination task for objects showing an example of a trial in the low (left) and high (right) feature ambiguity condition, respectively. C. Perceptual discrimination task for scenes showing an example of a trial in the low (left) and high (right) feature ambiguity condition, respectively. Red circles were not present in the actual task and are used here to illustrate the difference between stimulus features was different in the respective trial. This figure is adapted from the article Gellersen et al. (2021) published in *Cognition* (https://doi.org/10.1016/j.cognition.2020.104556).

**Figure 2.**
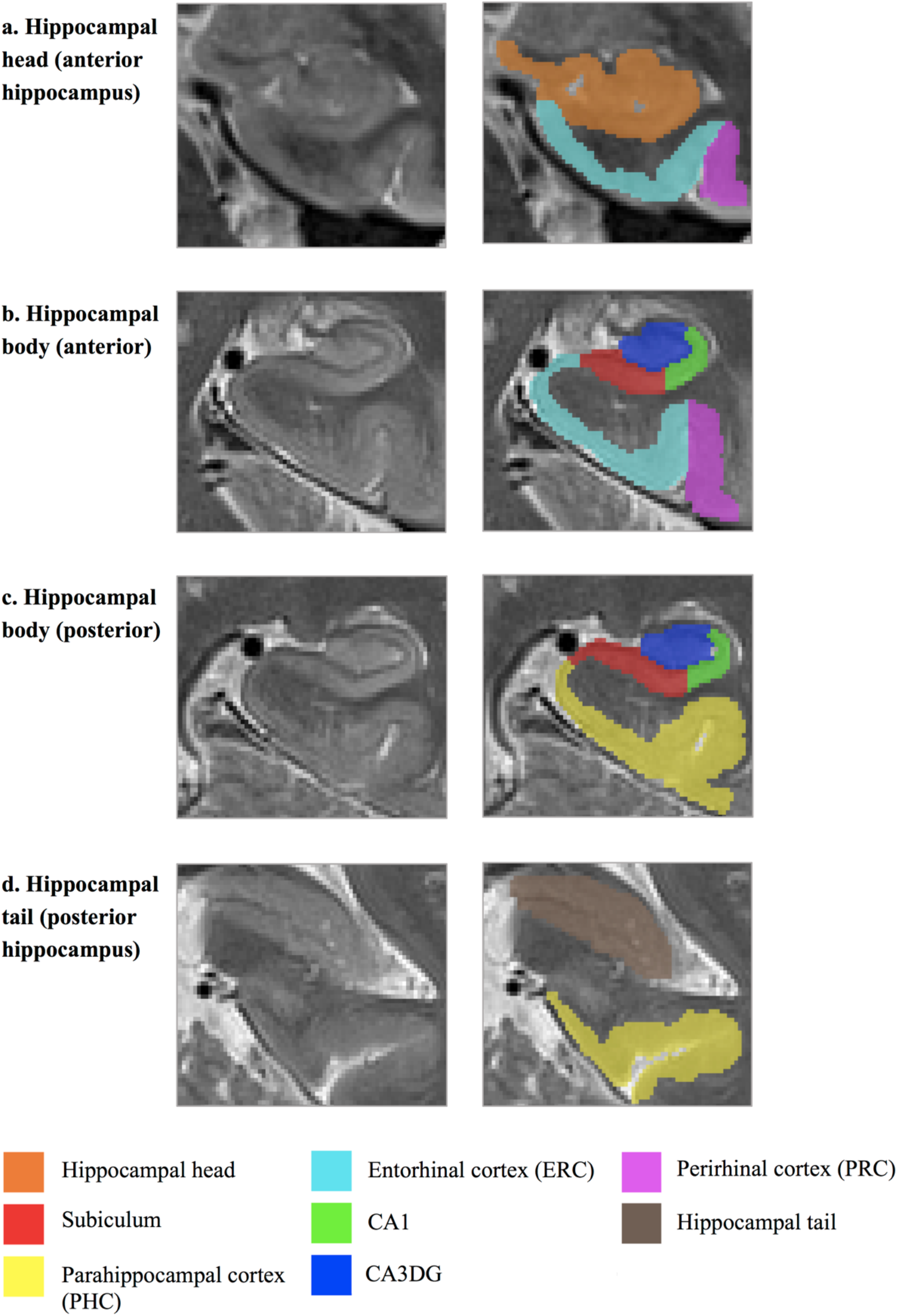
Example of a manual segmentation on a high resolution T2-weighted image.

The perceptual discrimination tasks were adapted from Barense, Gaffan & Graham (2007) for objects (“Greebles”) and Lee et al. (2005) for scenes. We previously used these tasks in the sample of cognitively normal older adults (Gellersen, Trelle, et al., 2021). Participants were asked to select the odd one out of three exemplars of either objects or scenes. Items either shared few perceptual features (low ambiguity condition) or were highly similar to one another and presented from different viewpoints (high ambiguity condition). As a result, in the low ambiguity condition items could be differentiated on the basis of basic perceptual features and in the high ambiguity task viewpoint-invariant processing of feature configurations was essential for discrimination. The order in which participants completed the two tests was counterbalanced across participants.

Stimuli for all tasks were presented in MATLAB *(Mathworks*, Inc., USA) in *Cogent 2000* (Cogent 2000 team at the FIL and the ICN and Cogent Graphics by John Romaya at the LON at the Wellcome Department of Imaging Neuroscience).

### Neuropsychological tests

Crystallised IQ was measured with the Vocabulary test of the Shipley Institute of Living Scale (Shipley, 1986) and the National Adult Reading Test (Nelson & Willison, 1991). Executive functioning and attention were tested using the Digit Span Forward and Backward from the Wechsler Adult Intelligence Scale (Wechsler, 2008) and Trails A and B and Verbal Fluency tests from the D-KEFS (Delis et al., 2001)._Finally, the Memory Functioning Questionnaire (MFQ) was administered to characterise subjective memory experience such as concerns about memory performance in daily life (Gilewski et al., 1990). We analysed data from two subscales of the MFQ: the frequency (32 items) and seriousness of forgetting (18 items). Responses varied on a 7-point Likert scale with low scores indicating more severe memory problems and higher scores representing no problems. We created an executive functioning composite score by taking the average *z*-score across the total digit span, Trails B and verbal fluency tests. See Table 1 for an overview of the neuropsychological profile of at-risk older adults compared to cognitively unimpaired participants.

### Magnetic resonance imaging sequences

For each participant, two structural images were taken on a 3T Siemens Trio MRI system (Erlangen, Germany) with a 32-channel head coil: 1) a whole-brain T1-weighted (1 × 1 × 1 mm) magnetisation-prepared rapid gradient-echo (MPRAGE) sequence with a repetition time (TR) of 2300ms, an echo time (TE) of 2.96ms, a field of view (FOV) of 256mm, flip angle of 9°, and 176 sagittal slices; and 2) a high-resolution T2-weighted turbo spin echo (TSE) image with TR/TE=8020/50ms, FOV=175mm and flip angle of 122° with partial brain coverage of 30 oblique coronal slices covering the medial temporal lobe at a resolution of 0.4 × 0.4 × 2 mm.

Eight T2-scans were excluded from all analyses with manually segmented volumes due to substantial motion artefacts. Another participant was excluded due to motion visible in the T1 scan. Finally, one participant was excluded from analyses on hippocampal volumes because of significant blurring that was restricted to this region. As a result, perirhinal and entorhinal volumes were available for 102 participants for manual segmentations (21 AR, 81 CU), whereas for the hippocampus this was the case for 91 individuals (21 AR, 70 CU) due to a technical error that cut off the hippocampal tail in eleven T2-scans. For ERC volumes and thickness derived from Freesurfer version 6, 106 scans were available (22 AR, 84 CU).

### Freesurfer segmentation

T1-weighted scans were used to obtain total intracranial volume (TIV) and MTL cortical thickness estimates based on the Desikan-Kiliany atlas (Desikan et al., 2006; Fischl & Dale, 2000). Thickness measures were extracted for entorhinal cortex and parahippocampal gyrus which in the Freesurfer parcellation includes both PRC and PHC. We calculated the average thickness across the two hemispheres. Total frontal volumes were obtained from the Freesurfer parcellation by summing volumes from left and right hemispheres of all regions defined to be located in the frontal lobes as described in https://surfer.nmr.mgh.harvard.edu/fswiki/CorticalParcellation.

### Manual Segmentation

MTL subregions were defined by manual segmentations of the T2 images using ITK-Snap software (Version 3.6.0; www.itksnap.org) (Yushkevich et al., 2006). Segmentations delineated perirhinal cortex (PRC), entorhinal cortex (ERC), parahippocampal cortex (PHC), and the hippocampus. Within the body of the hippocampus (beginning from the first slice after the disappearance of the uncus), hippocampal subfields CA1, subiculum, and a combined CA3/Dentate gyrus region were also delineated. Segmentations were conducted following a protocol developed by Carr and colleagues (2017). Manual segmentations of 103 T2-scans were randomly assigned to one of two raters (HMG; BGF) blinded to cognitive status. Prior to segmentation, the raters jointly coded the slice number of the onset and offset of each subregion and resolved any ambiguities in collateral sulcus type (shallow, normal, deep, double) which dictated the delineation of the PRC-ERC boundary. Six images were segmented by both raters in order to assess inter-rater reliability based on the Dice coefficient. The Dice metric indicated good to excellent inter-rater reliability ranging from .86 to .90 depending on the region with a mean of .89±.03. For all analyses, volumes of the two hemispheres were pooled.

Subregional volumes were corrected for TIV before analysis according to the formula Volume_corr_ = Volume_raw_ - β*(TIV_indiv_-TIV_mean_), where *β* refers to the regression coefficient of the model on a given regional volume of interest while using TIV as predictor. We chose this normalization method to maximize comparability with prior studies (Bennett et al., 2019; Doxey & Kirwan, 2015; K. L. Kern et al., 2021; Olsen et al., 2017; Stark & Stark, 2017).

### Statistical analysis

All analyses were carried out in RStudio version 1.2.5001.

In accordance with prior work on comparisons of Forced Choice and Yes/No performance (Bayley et al., 2008; Gellersen, Trelle, et al., 2021; Trelle et al., 2017), we calculated the discriminability index *d’* as follows:

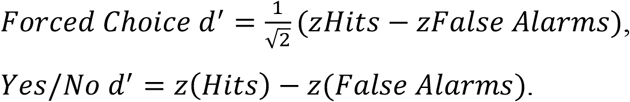

Accuracy of perceptual discrimination tasks was also transformed to *d*’ scores according to a three-alternative forced choice task. If the proportion of hits or misses was 0 or 1, we applied a correction to avoid a *d*’ of infinity: 0% misses were recoded as 1/(2*N*)=.005 and 100% hits as 1–1/(2*N*)=.995 with *N*=100 to reflect the number of trials (Macmillan & Creelman, 1991).

### MoCA group differences in mnemonic and perceptual discrimination

We excluded four participants from analyses of memory (*n*=1 AR) and perception (*n*=2 AR, *n*=1 CU) scores, respectively, because in three cases their performance suggested that they misunderstood the task instructions or were responding randomly (these data points are not excluded from Figure 3) and in one case there was a technical error during the acquisition of the perceptual discrimination task.

**Figure 3.**
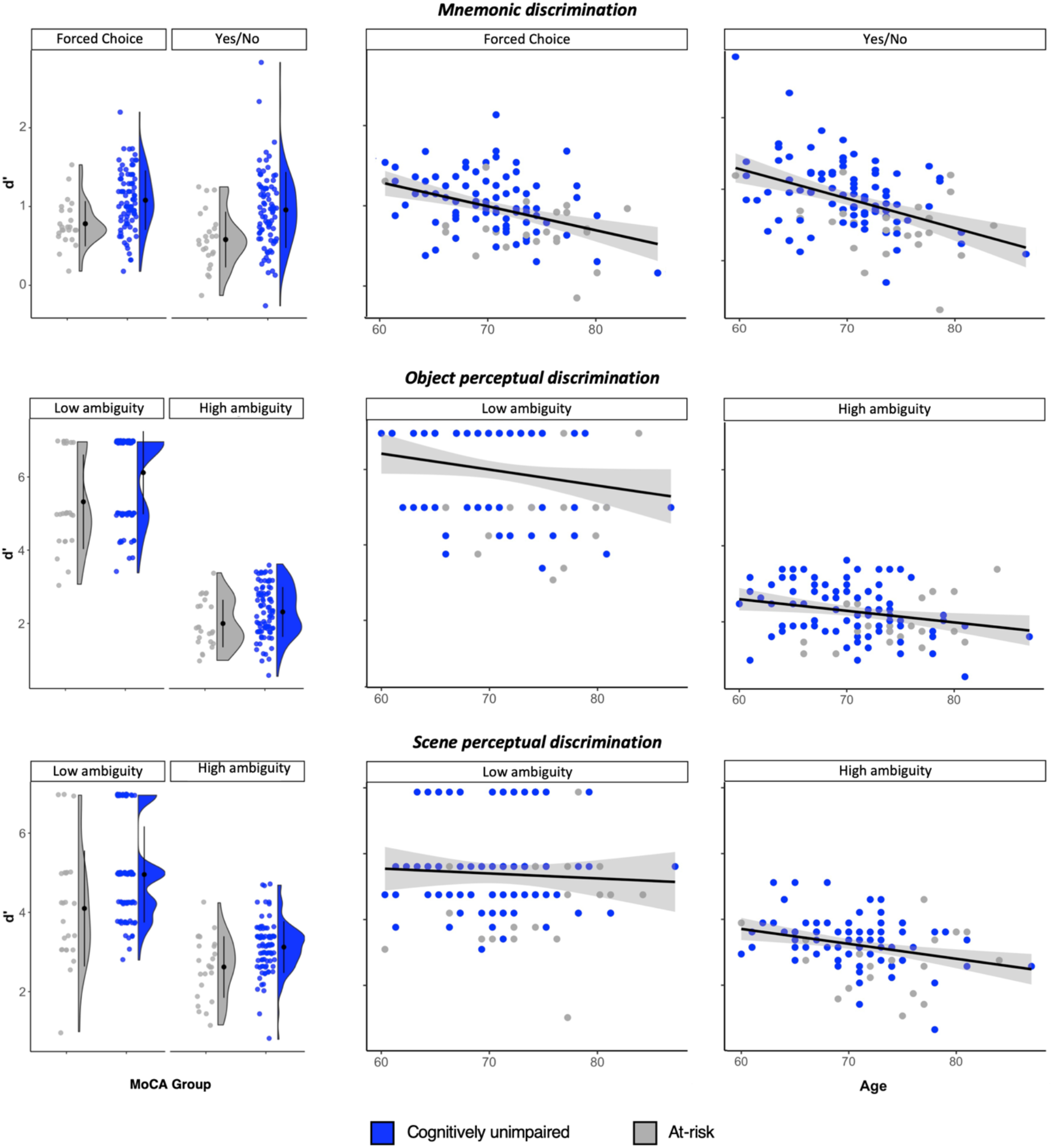
Effects of cognitive status and age on perceptual and mnemonic discrimination performance.

We used linear mixed-effects models with the *lmer* function from the *lme4* package in R (Bates et al., 2015) and tested whether cognitive status (cognitively unimpaired, at-risk for MCI) as indexed by MoCA scores was related to performance in perceptual and mnemonic discrimination. The dependent variable for the first model was performance on mnemonic discrimination tasks as indexed with *d*’. We compared candidate models in a stepwise fashion with the full model of interest being: *d*’ ~ *Age + Education + MoCA status (CU, AR) + Task Format (Forced Choice, Yes/No) + Age × Task Format × MoCA status + Error (by Subject ID)*, where individual participants were modelled as random intercepts. Based on our prior findings of age-related deficits in mnemonic discrimination being equivalent across the two tests, we did not include an age by task format interaction or an age by MoCA by task format interaction (Gellersen, Trelle, et al., 2021; Trelle et al., 2017).

We previously found that age effects were more pronounced for high than low feature ambiguity (Gellersen, Trelle, et al., 2021) and hypothesised that older adults at risk for MCI would show the greatest deficit in high-ambiguity object perceptual discrimination compared to CU older adults. For the model on perceptual discrimination scores, we therefore tested the full model with the highest-order interaction being one of age, MoCA status, ambiguity and category: *d’ ~ Education + Age × Ambiguity × Category × MoCA status + Error (by Subject ID)*, with all lower-order interactions included except for those with education (not shown here for the sake of brevity). Models were fitted using restricted maximum likelihood estimation. Post-hoc tests were carried out on estimated marginal means for a given effect of interest while controlling for multiple comparisons using Tukey correction. To determine which effects should be retained in the model, we used *F*-tests, Akaike Information Criterion (*AIC*), Bayesian Information Criterion *(BIC)* and Root Mean Squared Error *(RMSE)*.

Normality of residuals and homoskedasticity were assessed using Shapiro-Wilk’s test and Breusch-Pagan test, respectively, and visual inspection of diagnostic plots. Models were examined for extreme standardised residuals (±3) and for influential cases based on Cook’s distance with a cut-off of 4/number of cases in the model. Robustness tests for problematic cases involved case deletion and calculation of robust standard errors with the *robustlmm* package for mixed models (Koller, 2016), to determine whether a correction would result in a substantial change in the model.

The model on mnemonic discrimination scores violated the assumption of homoskedasticity. To account for the differences in variance of residuals of the two MoCA groups, we adjusted the variance-covariance matrix to allow for heterogeneity of variance on the basis of MoCA status. The model also had non-normally distributed residuals, but after removal of influential cases, the results remained robust and residuals were normally distributed. Given that the model effects remained robust to the removal of these cases, the non-normality of residuals did not change the interpretation of the model and we refrained from transforming the data to improve interpretability of the model.

### Mediation analysis: what factors can account for the effect of cognitive status on mnemonic discrimination?

We previously found that individual differences in Forced Choice and Yes/No mnemonic discrimination performance were differentially related to other cognitive abilities (Gellersen, Trelle, et al., 2021). Although perceptual discrimination correlated with mnemonic discrimination across both tasks, when comparing the relative contribution to performance in the two test formats, perceptual discrimination was more strongly associated with Forced Choice performance, while executive function better explained inter-individual differences in Yes/No performance. Here we followed up on our findings of mnemonic discrimination impairments in at-risk older adults by determining whether perceptual discrimination and executive functioning could account for the effect of cognitive status on memory as a function of task demands. Based on our finding from the mixed linear models which showed no statistical differences in the effects of MoCA status as a function of perceptual discrimination task stimulus category (objects, scenes) and feature ambiguity (low, high; see Results), we used a composite score of perceptual discrimination as predictor. Given the shared representational demands for any task requiring the disambiguation of perceptually highly similar stimuli and in order to optimise for statistical power, we conducted a mediation analysis with average mnemonic discrimination scores across tasks to test whether the total perceptual discrimination score could account for the relationship between MoCA group status and memory performance. We also conducted a mediation analysis to test whether executive deficits could account for the effect of MoCA status on Yes/No mnemonic discrimination. The MoCA contained shorter, simpler versions of the tasks used in our executive functioning composite (trail making, fluency and digit span). However, aside from a weak correlation between the respective digit span tasks (*r*=.26, *p*=.01), scores on the other tests were unrelated (*p*>.3). We therefore deemed it appropriate to conduct our analysis on cognitive status based on the total MoCA scores. The mediation analysis was carried out using ordinary least squares regression with the *mediation* package in R (Tingley et al., 2014) and statistical significance tests were based on bootstrapping with 10,000 iterations. Age was used as a covariate in all models.

### MTL subregional volumes and associations with cognition

Our structural measures of interest included total hippocampal, CA3DG, ERC, PRC and PHC volume, as well as ERC and PHC thickness. Associations between grey matter integrity and MCI risk status for these regions were probed using linear regressions. In the case of the model for ERC thickness, we adjusted the variance-covariance matrix using an HC3 estimator in the R *sandwich* package to mitigate the effect of heteroskedasticity in the model residuals (Zeileis, 2004; Zeileis et al., 2020). All analyses included age and education as covariates and were conducted on TIV-corrected volumes. Thickness measures were not normalised for TIV but corrected for sex (Westman et al., 2013). We then followed a stepwise procedure informed by best subset regression to probe for the contribution of different grey matter metrics to explaining variability in task performance. In this stepwise procedure, we selected models based on *AIC* with the criterion for model selection defined as *ΔAIC≤-2* in line with prior work in the field (Palmqvist et al., 2021). We tested for positive associations based on one-tailed tests of model coefficients. For the model on scene perception, we considered ERC, hippocampus and PHC measures as predictors. For object perception we included ERC and PRC. For the two mnemonic discrimination test formats, candidate predictors were ERC, PRC, CA3DG and total hippocampal grey matter metrics.

## Results

### Neuropsychological profile of cognitively unimpaired vs. at-risk older adults

Controlling for age, the at-risk group did not report significantly more instances of forgetting in their everyday lives (*F*(1, 105)=.98, *p*=.726), nor did they rate instances of forgetting as significantly more serious compared to the unimpaired group (*F*(1, 105)=1.95, *p*=.581). While the groups were defined by overall MoCA score, it is noteworthy that the at-risk group performed worse on all sub-scales of the test: attention, visuospatial/executive, language and memory, with the most apparent effect being on the memory subscale (all *p*<.001, controlled for age). The high-risk group also had lower scores on Shipley Vocabulary and the National Adult Reading tests, Trail Making test (version B) and the Digit Span test (all *p*<.001 while controlling for age).

### Older adults at risk for MCI are impaired in mnemonic and perceptual discrimination

A summary of mean scores on the memory and perception tasks can be found in Table 2. Figure 3 shows the effect of MoCA group status on mnemonic and perceptual discrimination performance (*d*’).

**Table 2.**
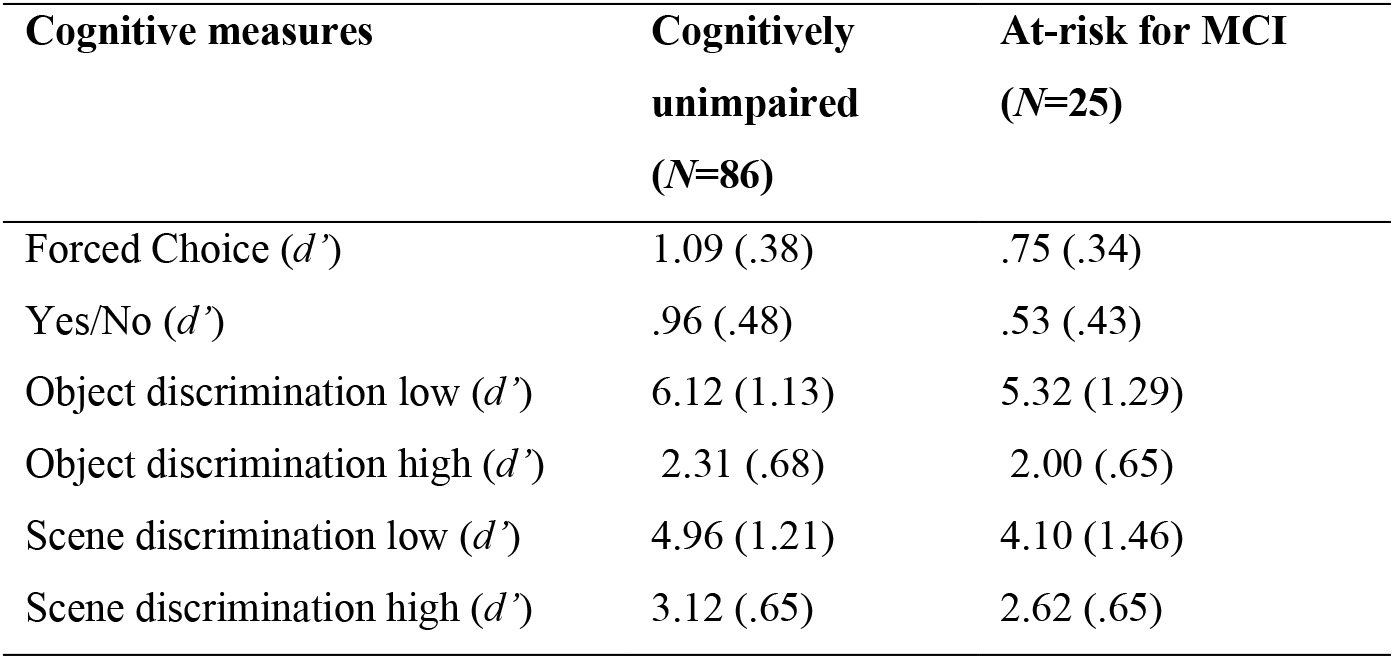
Performance on mnemonic and perceptual discrimination tasks by MoCA status.

| Cognitive measures | Cognitively<br>unimpaired<br>(N=86) | At-risk for MCI<br>(N=25) |
| --- | --- | --- |
| Forced Choice ($d'$) | 1.09 (.38) | .75 (.34) |
| Yes/No ($d'$) | .96 (.48) | .53 (.43) |
| Object discrimination low ($d'$) | 6.12 (1.13) | 5.32 (1.29) |
| Object discrimination high ($d'$) | 2.31 (.68) | 2.00 (.65) |
| Scene discrimination low ($d'$) | 4.96 (1.21) | 4.10 (1.46) |
| Scene discrimination high ($d'$) | 3.12 (.65) | 2.62 (.65) |

Mnemonic discrimination performance could best be explained based on main effects of format and MoCA status in addition to the covariates of age and education. The at-risk group performed significantly worse across mnemonic discrimination tasks relative to the unimpaired group (*F*(1, 103)=8.62, *β*=.530, *SE*=.180, *p*=.004). Older age was also associated with worse memory performance across test formats (*F*(1, 103)=24.56, *β*=-.295, *SE*=.075, *p*<.001). Across groups, performance was higher in the Forced Choice test format than the Yes/No test format (*F*(1, 106)=13.69, *β*=-.320, *SE*=.087, *p*<.001). Adding an interaction of MoCA group and task format did not improve the model (*ΔAIC*=2.58; interaction effect: *F*(1,105)=.73, *β*=.177, *SE*=.208, *p*=.396), suggesting that deficits in the two mnemonic discrimination tasks were equivalent in at-risk individuals. In a follow-up analysis we tested for the proportion of variance shared between the two tasks. Based on Stark and colleagues (2015) test-retest reliability for the mnemonic similarity task is .63. If the two tasks (FC, YN) were redundant, one should expect a similar *R^2^* value. However, our findings suggest that the two tasks capture different aspects of long-term memory based on an *R^2^* of half the expected value (23% and 36% of shared variance with and without controlling for age, respectively), based on our prior findings (Gellersen, Trelle, et al., 2021) and on the mediation analyses reported below.

Perceptual discrimination performance could best be accounted for using a model with age, education, stimulus category, feature ambiguity and MoCA status as main effects, and a two-way interaction between category and ambiguity. The at-risk group performed significantly worse in perceptual discrimination relative to the unimpaired group (*F*(101)=8.75, *β*=.269, *SE*=.091, *p*=.004). The magnitude of this impairment did not differ by stimulus category or feature ambiguity as the addition of either interaction effect did not improve the model (for MoCA by category: *ΔAIC*=3.15; interaction: *F*(1,311)=.23, *β*=.095, *SE*=.201, *p*=.475; for MoCA by ambiguity: *ΔAIC*=1.86; interaction: *F*(1,311)=1.52, *β*=.247, *SE*=.200, *p*=.219). Performance did not significantly vary as a function of age once MoCA status was included in the model (*F*(101)=2.58, *β*=-.060, *SE*=.037, *p*=.111). A stimulus category (object, scene) by feature ambiguity (high, low) interaction was also observed (*F*(312)=31.84, *β*=-1.101, *SE*=.092, *p*<.001), reflecting higher performance for scene stimuli relative to the object stimuli in the high ambiguity condition (*t*(312)=-6.83, *p*<.001), but higher performance on the object stimuli than the scene stimuli in the low ambiguity condition (*t*(312)=10.07, *p*<.001). Put another way, regardless of risk status, the effect of increasing feature ambiguity was more detrimental to performance for the object than the scene discrimination task (*t*(312)=-11.95, *p*<.001).

All results were robust to the removal of outliers and cases identified as having extreme residuals or greater influence on the regression coefficients. In a sensitivity analysis controlling for both education and premorbid IQ, as defined by the NART, these results remained robust. Given the potential role of working memory in perceptual oddity tasks, we conducted a sensitivity analysis controlling for executive functions which were added to the best fit model identified for perceptual discrimination. Despite an association between executive functions and perceptual discrimination (*F*(1,105)=4.94, *β*=.087, *SE*=.039, *p*=.028), the effect of risk status remained significant (*F*(1,107)=4.85, *β*=.212, *SE*=.096, *p*=.030).

### Factors accounting for mnemonic discrimination deficits

We next set out to determine whether different cognitive factors could explain the relationship between risk status and mnemonic discrimination. Figures 4A and 4B show the pairwise associations between executive functions, perceptual discrimination and mnemonic discrimination as a function of memory task format. Our analyses follow from our previous work in cognitively unimpaired older adults, which demonstrated that executive functions (a proxy for prefrontally-driven strategic retrieval abilities) and perceptual discrimination (a proxy for representational quality) made different contributions to individual differences in mnemonic discrimination performance as a function of task format (Gellersen, Trelle, et al., 2021), with executive functioning being relatively more important for Yes/No as opposed to Forced Choice performance due to the absence of retrieval support and the resulting need for strategic retrieval strategies such as recall-to-reject (Migo et al., 2009). To ensure that the same effects could still be observed when including the at-risk group in our sample we conducted stepwise regressions which reproduced the findings from Gellersen et al. (2021), finding perceptual discrimination as predictors in a model for Forced Choice and Forced Choice and executive functions as predictors of Yes/No performance (Supplementary Material). To provide a direct statistical comparison of the contribution of these two factors as a function of task format, we ran a mixed linear model with mnemonic discrimination scores as dependent variable and format (FC, YN) and cognitive process (EF, PD) as predictors. We found support for an interaction between these factors (*F*(1,294)=5.42, *β*=-.130, *SE*=.056, *p*=.021), with a post-hoc analysis showing that executive functions were indeed a significantly better predictor for Yes/No than Forced Choice performance *(Estimate=-.171, SE*=.046, *t*(296)=-3.74 *p*=.001). For the contribution of perceptual to mnemonic discrimination, there was no statistically significant difference between Forced Choice and Yes/No tasks (*Estimate*=-.041, *SE*=.032, *t*(296)=-1.27 *p*=.586).

**Figure 4.**
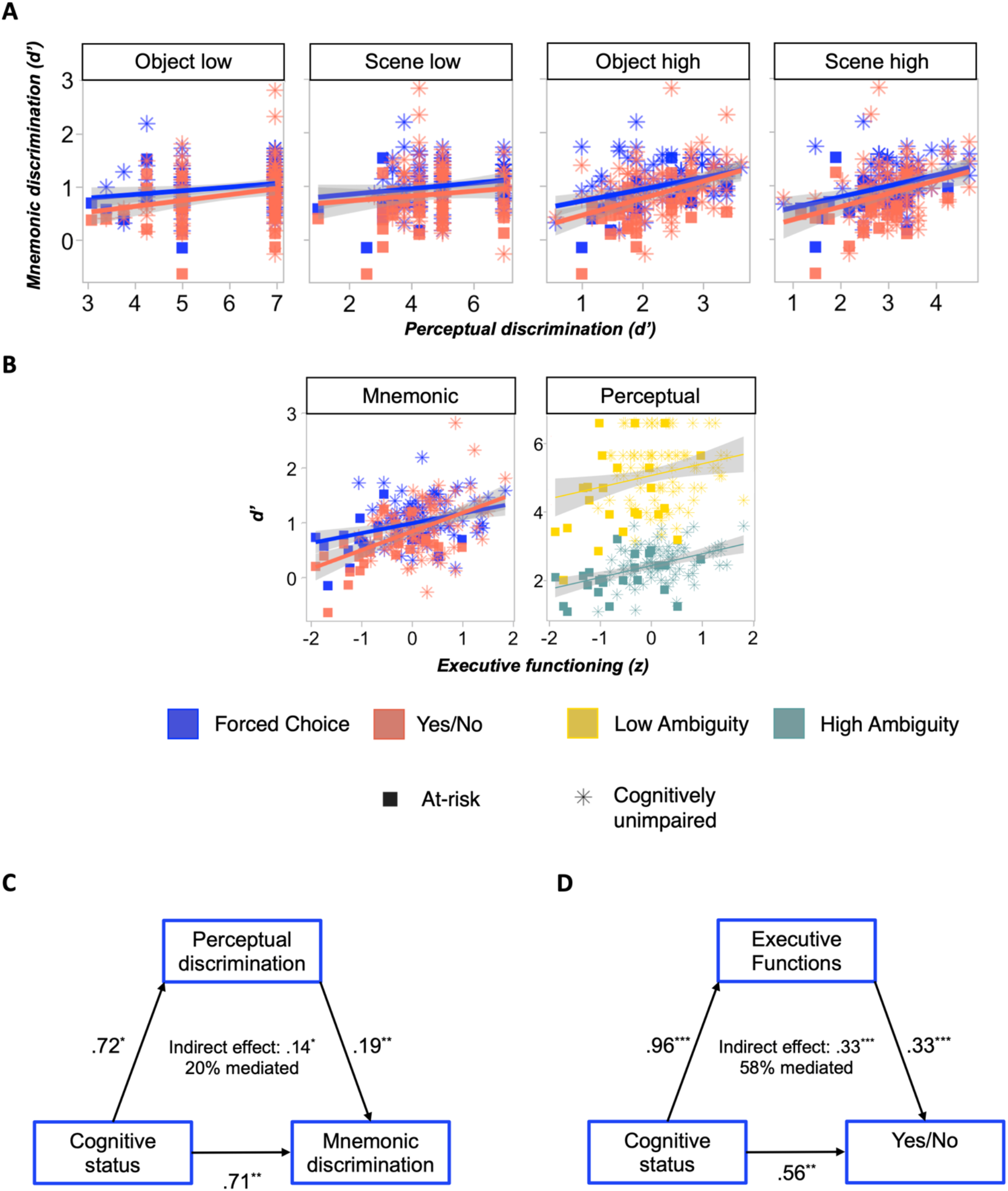
A. Associations between perceptual and mnemonic discrimination. B. Associations between executive functioning and perceptual and mnemonic discrimination, respectively. C. Mediation analysis with perceptual discrimination as mediator for the relationship between MoCA group and mnemonic discrimination total scores across formats. D. Mediation analysis with executive functioning as mediator for the relationship between MoCA group and mnemonic discrimination in the Yes/No task.

Using mediation analyses we next tested whether risk status reflects contributions from perceptual processes (generally important for mnemonic discrimination performance) and executive functions (additionally important for Yes/No) by testing each factor as a mediator of the relationship between MoCA group status and mnemonic discrimination. The mediation analysis on mean mnemonic discrimination scores across tasks (Figure 4C) showed a significant indirect effect of MoCA status via perceptual discrimination performance (*β*=.139, 95% *CI* [.001, .342], *p*=.048) which did not entirely account for the direct effect (*β*=.566, *95*% *CI* [.166, .971], *p*=.006). This suggests a small, partial mediation effect and a potential role for complex perceptual processes in mnemonic discrimination deficits in individuals at risk for cognitive decline.

Figure 4D shows the results of the mediation analysis for predicting Yes/No performance from MoCA group with neuropsychological tests of executive functions as a mediator. The total effect of MoCA status on Yes/No mnemonic discrimination (*β*=.558, 95% *CI* [.204, .939], *p*=.005) could be accounted for by an indirect effect via executive functioning as mediator (*β*=.326, *95% CI* [.113, .630], *p*<.001), rendering the direct effect of MoCA group on Yes/No performance non-significant (*β*=.232, *95% CI* [-.190, .623], *p*=.255), i.e, consistent with full mediation. A sensitivity analysis for Forced Choice performance demonstrated that executive dysfunction was not a mediator for poorer target-lure discriminability in at-risk older adults in this task with lower demands on strategic retrieval (*β*=.153, *95% CI* [-.123, .824], *p*=.262). A second sensitivity analysis showed that both Forced Choice (*β*=.462, *95% CI* [.300, .627], *p*<.001) and executive functioning (*β*=.300, *95% CI* [.132, .461], *p*<.001) made independent contributions to inter-individual variability in Yes/No performance. These sensitivity analyses dovetail with the results of the mediation analysis which support the notion that Yes/No performance additionally relies on cognitive control abilities, whereas Forced Choice deficits in the AR group cannot be explained by this factor.

### MoCA status is associated with entorhinal cortical thickness and volume

A summary of the relationship between MoCA group and MTL volumes and cortical thickness of our regions of interest can be found in Figure 5. Our analyses focused on the categorical MoCA group status. Individuals who failed the MoCA exhibited significantly reduced entorhinal cortical thickness (derived from Freesurfer: *β*=.741, *SE*=.332, *t*=2.23, *p*=.028, adjusted *R^2^*=.12, *f*^2^=.09 in a model with age and education as covariates) and volume (derived from manual segmentations: *β*=.533, *SE*=.249, *t*=2.14, *p*=.035, adjusted *R^2^*=.04, *f*^2^=.05). There were no differences between MoCA groups in terms of hippocampal volume (*p*=.125). For all other MTL subregions, no associations between MoCA status and grey matter were found (all *p*>.05).

**Figure 5.**
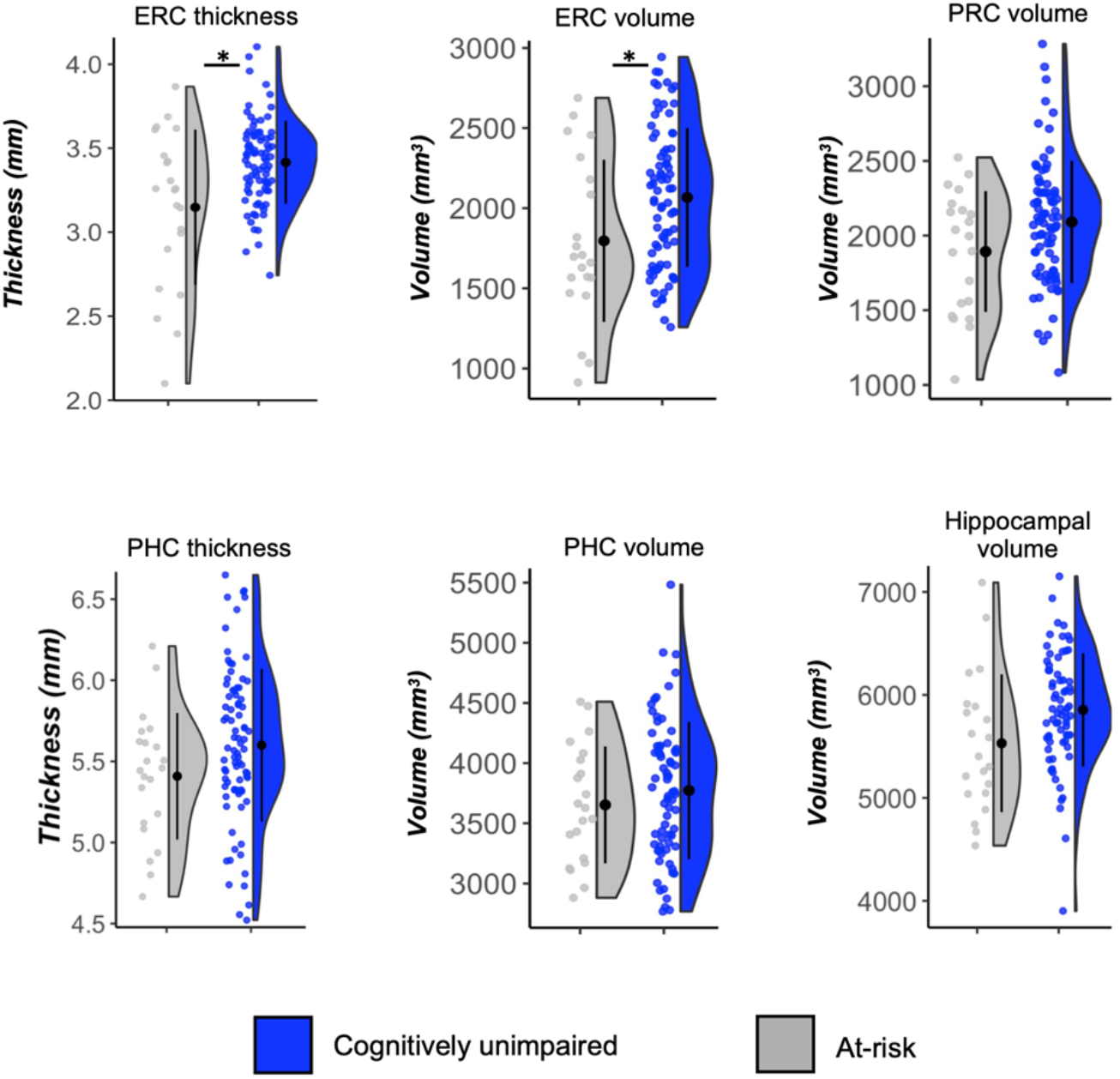
Medial temporal lobe subregional volumes and thickness by MoCA status. \**p*<.05

In an exploratory analysis, we also tested whether the at-risk group had lower frontal cortex volume given their pronounced executive functioning deficits. No differences between risk groups were found (*p*>.9).

### Associations between brain structure and cognition

Figure 6A presents brain-behaviour relationships for Yes/No or Forced Choice mnemonic discrimination. In no case did the addition of a grey matter measure for either of the models on mnemonic discrimination metrics pass the criteria for model selection (*ΔAIC*<2). ERC thickness showed a weak association with scene discrimination performance (model: *F*(3, 82)=2.83, *p*=.043, adjusted *R^2^*=.06, *f*^2^=.03) that trended towards significance (*β*=.152, *SE*=.094, *p*=.056), but made negligible improvement to the model fit (*ΔAIC=.66* over a model with age and education). The same was true for ERC thickness as a predictor in the object discrimination task (model: *F*(3, 83)=3.96, *p*=.011, adjusted *R^2^*=.09, *f*^2^=.05; ΔAIC=1.87 over a model with age and education; coefficient: *β*=.191, *SE*=.098, *p*=.028). Using a combined score of object and scene tasks in an exploratory analysis increased the effect size for ERC thickness, likely by reducing measurement error (*F*(3, 81)=4.33, *p*=.007, adjusted *R^2^*=.11, *f*^2^=.06; *ΔAIC*=3.04 over a model with age and education; ERC: *β*=.206, *SE*=.093, *p*=.029).

**Figure 6.**
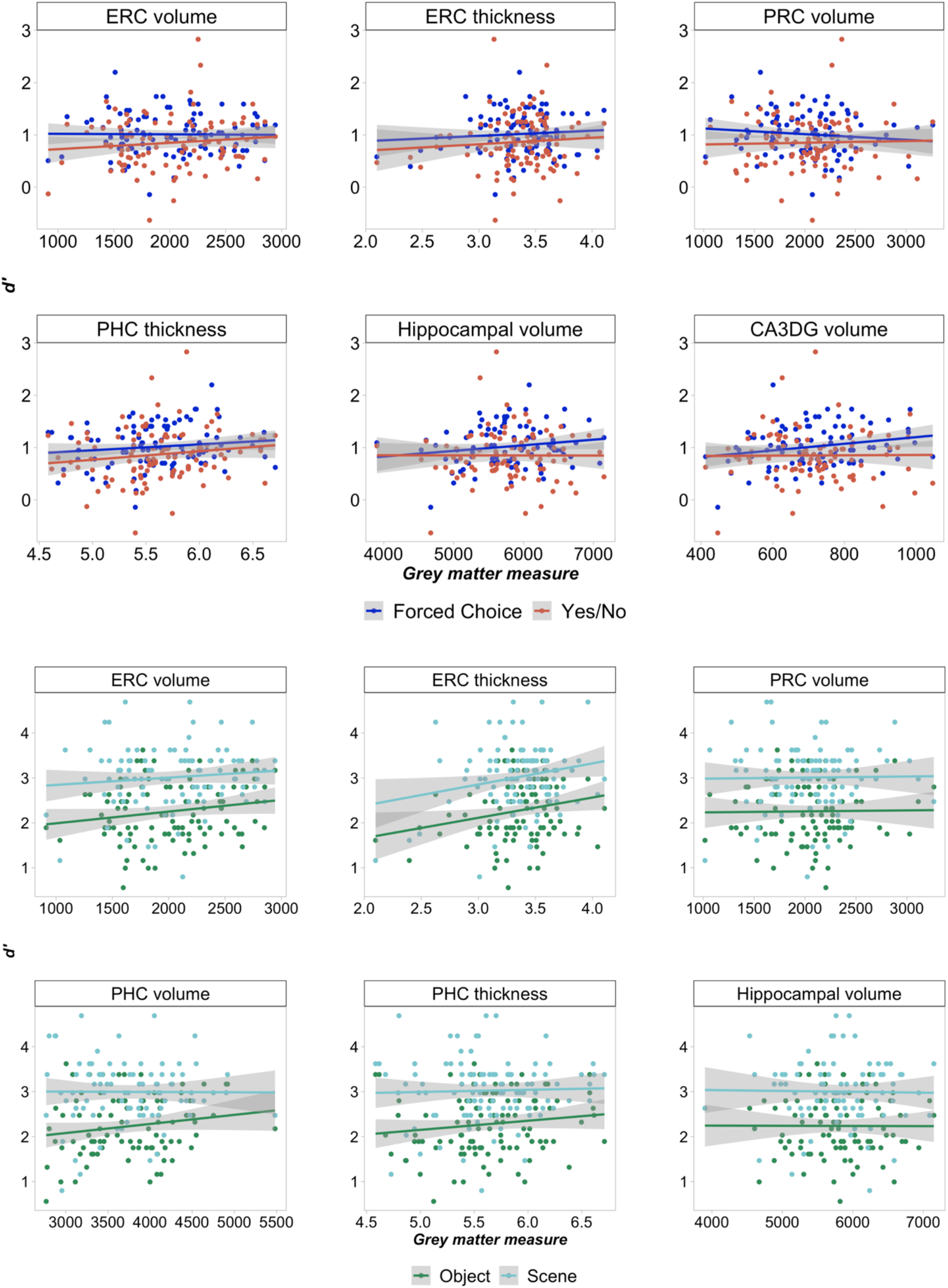
Associations between medial temporal lobe subregional integrity, perceptual and mnemonic discrimination. Note. Volumes are measured in mm^3^ and thickness in mm^2^. CA3DG: cornu ammonis 3 and dentate gyrus. ERC: entorhinal cortex. PHC: parahippocampal cortex. PRC: perirhinal cortex.

To test whether the association between performance on high-ambiguity perceptual discrimination and ERC thickness is greater than other structure-behaviour associations, we directly compared regression coefficients for entorhinal thickness and ERC, PRC, PHC and hippocampal volume. The effect for ERC thickness (*β*=.219, *95% CI* [.031, .407]) was not significantly different from that for ERC volume (*β*=.205, *95% CI* [-.028, .438]; *F*(1,77)=.01, *p*=.932), only marginally larger than the association with PRC (*β*=-.087, *95% CI* [-.312, .138]; *F*(1,77)=3.98,*p*=.050) and PHC volume (*β*=-.056, *95% CI* [-.265, .152]; *F*(1,77)=3.99, *p*=.050), and significantly larger than the association with hippocampal volume (*β*=-.279, *95% CI* [−.510, −.049]; *F*(1,77)=9.78, *p*=.002).

Follow-up exploratory analyses of brain-behaviour relationships for total frontal volumes revealed no association with executive functions or mnemonic discrimination (all *p*>.05). All brain-behaviour associations of interest as defined in the Methods section are shown in Figure 6.

## Discussion

In this study, we investigated cognitive and neural factors underpinning individual variability in mnemonic discrimination in healthy older adults and those at risk for cognitive impairment. We focused on older adults who have been identified as at risk for mild cognitive impairment based on a clinical screening tool (the Montreal Cognitive Assessment, MoCA) but who have not presented to a clinician. We found that mnemonic discrimination deficits in at-risk older adults were equivalent across Forced Choice and Yes/No test formats. At-risk individuals also demonstrated a deficit in perceptual discrimination of objects and scenes. This was not only when items were characterised by high feature overlap, but, contrary to our predictions, also when feature ambiguity was lowered. This deficit across all perceptual tasks mediated the relationship between cognitive risk status and mnemonic discrimination. In contrast, executive dysfunction accounted for Yes/No but not Forced Choice performance. At-risk older adults had reduced entorhinal cortical thickness and volume. Across all older adults, there was little evidence that MTL grey matter structure was related to inter-individual differences in mnemonic and perceptual discrimination performance, bar a small effect of entorhinal thickness on perceptual discrimination when combined across high ambiguity object and scene discrimination.

Our finding that mnemonic discrimination metrics are sensitive to cognitive risk status in older adults is in line with prior studies using Yes/No and Old/Similar/New paradigms (Bennett et al., 2019; Dillon et al., 2017; Pishdadian et al., 2020; Reagh et al., 2014; Trelle et al., 2021; Yassa et al., 2010). We further probed the nature of mnemonic discrimination deficits in at-risk older adults by contrasting Yes/No and Forced Choice formats to manipulate the degree to which mnemonic discrimination performance would place demands on PFC-mediated strategic retrieval processes and hippocampal-dependent mechanisms (Bastin & van der Linden, 2003; Gallo, 2004; Holdstock et al., 2002; Migo et al., 2009; Norman, 2010; Yonelinas, 2002). Previous studies have demonstrated that pattern separation deficits exist in older adults with memory impairment and MCI and contribute to mnemonic discrimination impairment in Old/Similar/New task formats (Reagh et al., 2014; Yassa et al., 2010). Whether these mnemonic deficits are equally pronounced in Forced Choice performance was less clear. Our finding of Forced Choice deficits in at-risk older adults suggests that reducing demands on recollection processes and allowing greater reliance on gist-based responding is not sufficient to rescue performance in this group (Bastin & van der Linden, 2003; Gallo, 2004; Gallo et al., 2006; Norman, 2010; Yonelinas, 2002). These results parallel those in prior work in community-dwelling older adults (Gellersen, Trelle, et al., 2021; Huffman & Stark, 2017; Trelle et al., 2017) and confirm our hypothesis that memory tasks reliant on high-fidelity representations formed by MTL subregions are sensitive to MCI risk even if recollection is not required for good performance.

To further this point, we also tested whether at-risk older adults would show comparable deficits in perceptual discrimination of objects and scenes under conditions of high feature ambiguity, task conditions in which demands on complex, conjunctive MTL-dependent representations are thought to be similarly high. Although it has previously been suggested that such tasks may be sensitive to risk for cognitive decline due to their reliance on MTL (Anderson, 2019; Fidalgo et al., 2016a; Grande et al., 2021), perceptual discrimination has rarely been tested empirically for this purpose. Our finding that older adults at risk for MCI are not only impaired on mnemonic but also on perceptual discrimination tasks lends support to these proposals. Interestingly, individuals at risk for MCI were also impaired on low ambiguity perceptual discrimination task. While such a deficit was not expected in a non-clinical population, this pattern suggests decline in representational quality beyond the MTL in ventral visual cortex (Lemos et al., 2016). Future work using AD biomarkers is necessary to provide direct evidence as to whether perceptual discriminations tasks are sensitive to early and subtle cognitive decline associated with preclinical AD.

Our finding that task demands may affect mnemonic discrimination deficits in older adults at risk for MCI is of particular interest for future work on early detection of memory decline using these tests. Although the magnitude of decline in mnemonic discrimination performance was equivalent across task formats among at-risk older adults, our results suggest that the interpretation of these deficits should be considered carefully, because task formats could have a major influence on the underlying processes that give rise to such deficits. Perceptual discrimination deficits may contribute to decline in mnemonic discrimination observed in high-risk individuals, both in our study and in previous work, regardless of task format (Berron et al., 2019; Sinha et al., 2018; Stark et al., 2013; Trelle et al., 2021; Webb et al., 2020; Yassa et al., 2010). This is likely because less detailed perceptual representations not only hamper hippocampal pattern separation but also weaken the distinctiveness of a cortical strength-based memory signal by virtue of leaving memory representations more vulnerable to interference (Kent et al., 2016; Ly et al., 2013; Monti et al., 2014; Muecke et al., 2018; Newsome et al., 2012; Ryan et al., 2012).

In contrast, the findings from our analysis with executive functions as mediator suggest that in mnemonic similarity tests with the Yes/No or Old/New/Similar format used in most studies to date, PFC-dependent processes, in addition to hippocampal recall, significantly contribute to the relationship between cognitive risk status and mnemonic discrimination performance (Adams et al., 2020; Bencze et al., 2021; Foster et al., 2020; Kirwan & Stark, 2007; Reagh et al., 2018; Sinha et al., 2018; Stark et al., 2019; Webb et al., 2020). In line with this proposal, Pishdadian et al. (2020) showed that performance in the traditional Old/New/Similar mnemonic similarity task was associated with total MoCA scores and scores on the executive functioning subscore but not with the memory subscale of the MoCA. These findings support the notion that this test format for mnemonic discrimination is sensitive to risk for general cognitive impairment but suggests that the task is not necessarily specific to indexing hippocampal-dependent changes to memory. Prefrontal atrophy and dysfunction is common in old age and in a variety of neurological conditions besides Alzheimer’s disease (Boyle et al., 2004; Lockwood et al., 2002) and entorhinal thinning tends to precede marked PFC atrophy as shown here and in prior work on preclinical AD (Mak et al., 2017). Reducing demands on executive functioning may therefore increase the specificity of mnemonic discrimination metrics for early detection of detrimental memory changes. Given that executive functions explained little overall variance in Forced Choice performance and could not account for performance deficits among the at-risk older adult group, our findings show that the Forced Choice test may provide valuable complementary information to performance probed with Yes/No or Old/New/Similar test versions. Future studies should explore the influence of mnemonic discrimination test format in the context of AD biomarkers to determine sensitivity to amyloid and tau burden in older adults prior to clinical decline.

Although the present study sought to isolate MTL-dependent perceptual discrimination ability using high ambiguity oddity tasks, it is not possible to rule out working memory contributions to task performance. However, we maintain that the most dominant characteristic impacting performance on this task are demands on complex feature conjunctions supported by the MTL, as demonstrated by a wealth of patient and neuroimaging work that have leveraged these stimuli previously (Barense et al., 2007, 2010; Erez et al., 2013; A. C. H. Lee et al., 2005). Moreover, our results show that associations between mnemonic and perceptual discrimination are not due to potential shared demands on cognitive control demands given that perceptual discrimination scores, but not executive functions, predicted individual differences in Forced Choice performance. Thus, any contributions of working memory processes to performance are unlikely to fully account for our findings of poorer complex perceptual discrimination in older adults at risk for MCI.

It is worth noting that while we and others have provided evidence for impaired perceptual discrimination in older adults, particularly under conditions of high feature ambiguity (Gellersen, Trelle, et al., 2021; Ryan et al., 2012; Trelle et al., 2017), this is not always observed. For example, a study by Yassa and colleagues (2011) did not find an age deficit in a perceptual/short-term memory version of the mnemonic discrimination task in which a studied object was followed by a brief noise mask after which a target, lure, or novel foil was presented. One key difference between this task and the perceptual discrimination task used here is that stimuli were presented from the same viewpoint as opposed to different angles. Prior work has demonstrated that the involvement of the MTL in perceptual discrimination tasks, and the likelihood that perceptual discrimination is impaired with age, is related to the degree a task places demands on viewpoint-invariant representations (Barense et al., 2007; Ryan et al., 2012). In contrast, when stimuli are presented from the same viewpoint, a single feature strategy can more easily disambiguate targets and foils and performance is less likely to rely on the MTL or be impaired in older adults. This strategy may be sufficient to maintain adequate performance in the short-term mnemonic discrimination task without feature interference. This strategy may become less reliable at higher memory load (Gellersen, 2022) and longer study-test delays where the ability to form highly detailed, unique stimulus representations may be relatively more important. These greater demands on viewpoint-invariant processing and the formation of detailed holistic representations robust to accumulating feature interference may explain why age effects were observed in our perceptual and long-term mnemonic discrimination tasks, respectively, but not in the study by Yassa and colleagues (2011). Thus, we do not see their results as incompatible with the current findings. However, they do highlight the complexity of exploring the role of the MTL in perception, including the key role of stimulus set features, as noted in recent computational work (Bonnen et al., 2021).

As expected, and in line with the literature, individuals with cognitive impairment exhibited reduced ERC thickness and volume relative to the unimpaired group (Devanand et al., 2008; Zhang et al., 2011). Structural changes in the ERC often correlate with tau deposition even in cognitively unimpaired older adults, raising the possibility that some proportion of individuals failing the MoCA may exhibit such pathology (Maass et al., 2018; Schroeder et al., 2017; Thomas et al., 2020). Regardless of underlying biomarker status, observed differences in ERC thickness suggest that the MoCA is sensitive to differences in MTL structural integrity among community-dwelling older adults without subjective memory complaints or a clinical diagnosis, as suggested by prior work (Olsen et al., 2017). Given that reduced MTL structural integrity is itself a key risk factor for cognitive decline (Holbrook et al., 2019), this finding lends support to the idea that MoCA underperformers are “at-risk” for cognitive impairment. Future work exploring risk for cognitive impairment among community-dwelling volunteers would benefit from a full neuropsychological evaluation, as well as the inclusion of in vivo AD biomarkers.

We expected that the use of a gold standard manual segmentation method and MTL-dependent sensitive memory and perception tasks would allow us to detect associations between perceptual-mnemonic processes and MTL subregional structure. However, we only found partial support for associations between brain structure and cognition. Individual differences in perceptual, but not mnemonic discrimination, were weakly associated with entorhinal cortical thickness. Given that ERC thickness was strongly related to MoCA status, our statistical power to detect a structural effect was likely greatest for ERC over other subregions. The absence of a detectable relationship between MTL subregions and memory scores may be surprising, as one would expect that variance in MTL structure would translate to the memory domain. However, the association between ERC thickness and perceptual discrimination was small and only held when combining scores from the object and scene tasks. We also found only weak support for this brain-behaviour relationship being specific to entorhinal thickness as opposed to other MTL grey matter measures. These data suggest that most of the associations between regional MTL grey matter measures and cognitive performance may have been too subtle as to be detected in the present sample, possibly due to statistical power and/or the fact that the majority of participants were cognitively normal. Future work in larger samples as well as measurement of changes in MTL structural integrity over time may also offer increased sensitivity compared to the present cross-sectional analysis, which cannot decouple natural variation in MTL subregion volume from within-subject atrophy.

It has previously been shown that entorhinal cortical thinning associated with AD pathology can be present without showing any correlation with memory decline (Doherty et al., 2015). Functional alterations such as hyperactivity in CA3 subfield and changes to the connectivity of subregions within MTL circuits are often an indicator of mnemonic discrimination deficits prior to the emergence of associations between structure and memory performance (Adams et al., 2020; Berron et al., 2019; Carr et al., 2017; Reagh et al., 2018; Sinha et al., 2018; Stark et al., 2020; Yassa et al., 2010, 2011). Our findings may provide boundary conditions for the relationship between brain macroscale anatomical features and mnemonic discrimination: in prior work on mnemonic discrimination that uncovered associations between MTL subregional grey matter structure and performance, studies often included both healthy older adults and diagnosed MCI cases (Bennett et al., 2019; Besson et al., 2020; Dillon et al., 2017), were enriched for the presence of preclinical AD pathology (Berron et al., 2020), or conducted their analysis across older and younger adults (Doxey & Kirwan, 2015; Stark & Stark, 2017). In all cases, variability in terms of memory performance and brain structure would be expected to be greater than in our study. Our data therefore suggest that brain behaviour-relationships are often elusive in older adults without clinical deficits, even when using tasks optimised to tap into the functions of specific MTL subregions.

This study has limitations that should be considered when interpreting the present findings. We did not obtain quantitative data pertaining to visual acuity which has previously been shown to contribute to mnemonic discrimination (Davidson et al., 2019). We can therefore not rule out that such basic perceptual factors made some contribution to interindividual variability in our tasks. However, it is unlikely that basic sensory decline is the main driver of the deficit in perceptual and mnemonic discrimination in at-risk older adults given that their cognitive status was defined on tasks not reliant on such properties of the visual system. Furthermore, the use of a cross-sectional design means that we could not measure intra-individual changes in cognitive ability over time. For example, some subjects may have lower baseline cognition, rather than exhibiting meaningful decline (e.g., due to aging/neurodegeneration; Kocagoncu et al., 2022). To control for effects of normal interindividual cognitive ability we relied on other proxies, controlling for education in all our analyses and further conducted sensitivity analyses controlling for premorbid intelligence. Neither factor changed the interpretation of our results.

Another limitation is the absence of biomarker data concerning potential AD pathology. Individuals with MCI or at risk for MCI are a heterogeneous group (Mariani et al., 2007) and not every adult identified as at-risk in our study will move on to develop clinical memory impairment or receive an AD diagnosis. Nonetheless, we argue that even in the absence of such biomarker data, our at-risk group is highly likely to harbour more individuals who will move on to develop MCI and potentially AD. First, prevalence of amyloid pathology in cognitively normal individuals aged 60+ years ranges between 10-30% (Chételat et al., 2013; Jansen et al., 2015; S. Kern et al., 2018), and is significantly higher in those with signs of cognitive impairment (Jansen et al., 2015). Second, older adults with a similar cognitive profile as our at-risk group are two times as likely to move on to develop AD as their cognitively normal counterparts (Parnetti et al., 2019). Third, entorhinal thinning is an independent predictor of AD conversion over and above cognitive status (Betthauser et al., 2020; Doraiswamy et al., 2012; Rowe et al., 2010; Sperling et al., 2013; Stoub et al., 2005) and the grey matter differences that were observed in the ERC in our at-risk group may provide an indirect marker of potential AD pathology given prior work showing associations between ERC thickness and tau pathology (Holbrook et al., 2019; Velayudhan et al., 2013). Moreover, our sample of older adults has similar characteristics (failing the MoCA, no MCI diagnosis, no cognitive complaints) compared to other at-risk groups identified in comparable studies which present findings of ERC volume loss (Olsen et al., 2017) and mnemonic discrimination impairment (Fidalgo et al., 2016b; Pishdadian et al., 2020; Yeung et al., 2013) that dovetail with the current results. Our findings can therefore still offer valuable insights into the cognitive profile of these at-risk older adults. Future work employing in vivo biomarkers is needed to provide further evidence in particular for the potential utility of complex perceptual discrimination tasks for the early detection of AD risk.

In conclusion, the present results add to a growing body of work demonstrating that individuals who perform below the normal range on the MoCA exhibit distinct structural and cognitive profiles from unimpaired older adults (Newsome et al., 2012; Olsen et al., 2017; Yeung et al., 2013, 2017, 2019), which includes deficits in both mnemonic and perceptual discrimination tasks and reduced cortical thickness and volume in ERC. Moreover, our results suggest that the interpretation of mnemonic discrimination deficit depends on task format, whereby executive functions mediate Yes/No performance, while perceptual discrimination, as proxy for representational quality, mediates mnemonic discrimination impairment across task formats. Finally, our findings suggest that MTL-dependent tasks, including both perceptual and mnemonic discrimination of stimuli with overlapping features, may be sensitive to risk for clinical cognitive impairment and should be explored in the context of future biomarker studies to better understand the potential utility of these tasks for early detection of Alzheimer’s disease and monitoring of cognitive change in older adults.

## Acknowledgements

Preliminary results of this study have been presented at the following conferences: Aging of Memory Functions, 2018, Bordeaux, France; Cognitive Neuroscience Society Annual Meeting, 2019, San Francisco, United States; Cambridge Memory Meeting, 2020, Cambridge, United Kingdom. Cognitive tasks have previously been described in Gellersen, H. M., Trelle, A. N., Henson, R. N., & Simons, J. S. (2021). Executive function and high ambiguity perceptual discrimination contribute to individual differences in mnemonic discrimination in older adults. *Cognition, 209*, 104556. Data from the cognitively normal older adult group in the current article belongs to a subset of the dataset presented the Gellersen et al. (2021) *Cognition* paper. This research was supported by the BBSRC [grant number BB/ L02263X/1], and was carried out within the University of Cambridge Behavioural and Clinical Neuroscience Institute, funded by a joint award from the Medical Research Council and the Wellcome Trust. H.M.G. was supported by a Medical Research Council doctoral training grant [#RG86932] and a Pinsent Darwin Studentship, B.G.F and S.M.K. by Biotechnology and Biological Sciences Research Council (BBSRC) doctoral training grants, R.N.H. by Medical Research Council programme grant [grant number SUAG/046 G101400], and J.S.S. by James S. McDonnell Foundation Scholar award [grant number #220020333]. The funders had no role in the conceptualisation, analysis or publication of this data. The study was conducted at the Behavioural and Clinical Neuroscience Institute (BCNI) and the MRC Cognition and Brain Sciences Unit (CBU) at the University of Cambridge. The authors would like to thank the funders for their support of their research. Special thanks go to Priyanga Jeyarathnarajah, Sarah Fox and Megan Thomson for help with data collection, to Mark Haggard for advice on data analysis, and to Morgan Barense for sharing task stimuli. For the purpose of open access, the author has applied a Creative Commons Attribution (CC BY) licence to any Author Accepted Manuscript version arising from this submission

## Data access statement

The behavioural data are available on the first author’s account on the Open Science Framework: osf.io/6suyd. Neuroimaging data are currently being analysed for further projects and can be made available via email request to the first author.

