## Supplementary Material for "Medial temporal lobe structure, mnemonic and perceptual discrimination in healthy older adults and those at risk for mild cognitive impairment"

ExecF = Executive functions
FC_dprime = Forced Choice performance
MDdprime = Combined mnemonic discrimination scores across task formats
MoCA = Montreal Cognitive Assessment
PD_all_dprime = Combined perceptual discrimination scores across all category and ambiguity levels
YN_dprime = Yes/No performance

1. Model selection for combined mnemonic discrimination performance

```
Forward Selection Method
------------------------

Candidate Terms:

1 . Age
2 . Education
3 . MoCA_Group
4 . PD_all_dprime
5 . ExecF

 Step 0: AIC = 298.1344
 MDdprime ~ 1
```

| Variable | DF | AIC | Sum Sq | RSS | R-Sq | Adj. R-Sq |
|---|---|---|---|---|---|---|
| ExecF | 1 | 274.122 | 22.793 | 80.207 | 0.221 | 0.214 |
| Age | 1 | 276.507 | 20.933 | 82.067 | 0.203 | 0.195 |
| MoCA_Group | 1 | 278.711 | 19.175 | 83.825 | 0.186 | 0.178 |
| PD_all_dprime | 1 | 282.985 | 15.658 | 87.342 | 0.152 | 0.144 |
| Education | 1 | 299.694 | 0.435 | 102.565 | 0.004 | -0.006 |

```
+ ExecF


 Step 1 : AIC = 274.122
 MDdprime ~ ExecF
```

| Variable | DF | AIC | Sum Sq | RSS | R-Sq | Adj. R-Sq |
|---|---|---|---|---|---|---|
| Age | 1 | 259.833 | 11.628 | 68.579 | 0.334 | 0.321 |
| MoCA_Group | 1 | 268.160 | 5.911 | 74.296 | 0.279 | 0.264 |
| PD_all_dprime | 1 | 268.460 | 5.697 | 74.510 | 0.277 | 0.262 |
| Education | 1 | 276.122 | 0.000 | 80.207 | 0.221 | 0.206 |

+ Age

Step 2 : AIC = 259.8326
MDdprime ~ ExecF + Age

| Variable | DF | AIC | Sum Sq | RSS | R-Sq | Adj. R-Sq |
|---|---|---|---|---|---|---|
| PD_all_dprime | 1 | 256.868 | 3.197 | 65.382 | 0.365 | 0.346 |
| MoCA_Group | 1 | 257.686 | 2.681 | 65.898 | 0.360 | 0.341 |
| Education | 1 | 261.826 | 0.004 | 68.574 | 0.334 | 0.314 |

+ PD_all_dprime

Step 3 : AIC = 256.8678
MDdprime ~ ExecF + Age + PD_all_dprime

| Variable | DF | AIC | Sum Sq | RSS | R-Sq | Adj. R-Sq |
|---|---|---|---|---|---|---|
| MoCA_Group | 1 | 256.170 | 1.674 | 63.708 | 0.381 | 0.356 |
| Education | 1 | 258.843 | 0.016 | 65.366 | 0.365 | 0.340 |

+ MoCA_Group

Step 4 : AIC = 256.1701
MDdprime ~ ExecF + Age + PD_all_dprime + MoCA_Group

| Variable | DF | AIC | Sum Sq | RSS | R-Sq | Adj. R-Sq |
|---|---|---|---|---|---|---|
| Education | 1 | 258.165 | 0.003 | 63.704 | 0.382 | 0.350 |

No more variables to be added.

Variables Entered:

+ ExecF
+ Age
+ PD_all_dprime
+ MoCA_Group

Final Model Output

### Model Summary

| | | | |
|---|---|---|---|
| R | 0.618 | RMSE | 0.802 |
| R-Squared | 0.381 | Coef. Var | 3155701964907629568.000 |
| Adj. R-Squared | 0.356 | MSE | 0.644 |
| Pred R-Squared | 0.313 | MAE | 0.594 |

RMSE: Root Mean Square Error
MSE: Mean Square Error
MAE: Mean Absolute Error

### ANOVA

| | Sum of Squares | DF | Mean Square | F | Sig. |
|---|---|---|---|---|---|
| Regression | 39.292 | 4 | 9.823 | 15.265 | 0.0000 |
| Residual | 63.708 | 99 | 0.644 | | |
| Total | 103.000 | 103 | | | |

### Parameter Estimates

| model | Beta | Std. Error | Std. Beta | t | Sig | lower | upper |
|---|---|---|---|---|---|---|---|
| (Intercept) | -0.284 | 0.193 | | -1.473 | 0.144 | -0.667 | 0.099 |
| ExecF | 0.263 | 0.092 | 0.263 | 2.848 | 0.005 | 0.080 | 0.446 |
| Age | -0.285 | 0.086 | -0.285 | -3.337 | 0.001 | -0.455 | -0.116 |
| PD_all_dprime | 0.163 | 0.088 | 0.163 | 1.845 | 0.068 | -0.012 | 0.338 |
| MoCA_GroupPass | 0.365 | 0.226 | 0.152 | 1.613 | 0.110 | -0.084 | 0.814 |

### Selection Summary

| Variable | AIC | Sum Sq | RSS | R-Sq | Adj. R-Sq |
|---|---|---|---|---|---|
| ExecF | 274.122 | 22.793 | 80.207 | 0.22129 | 0.21366 |
| Age | 259.833 | 34.421 | 68.579 | 0.33419 | 0.32100 |
| PD_all_dprime | 256.868 | 37.618 | 65.382 | 0.36523 | 0.34618 |
| MoCA_Group | 256.170 | 39.292 | 63.708 | 0.38148 | 0.35649 |

2. Model selection for Yes/No performance

```
Forward Selection Method
------------------------

Candidate Terms:

1 . Age
2 . Education
3 . MoCA_Group
4 . PD_all_dprime
5 . ExecF
6 . FC_dprime

 Step 0: AIC = 298.1344
 YN_dprime ~ 1
```

| Variable | DF | AIC | Sum Sq | RSS | R-Sq | Adj. R-Sq |
|---|---|---|---|---|---|---|
| FC_dprime | 1 | 246.237 | 41.657 | 61.343 | 0.404 | 0.399 |
| ExecF | 1 | 270.520 | 25.523 | 77.477 | 0.248 | 0.240 |
| Age | 1 | 279.299 | 18.700 | 84.300 | 0.182 | 0.174 |
| MoCA_Group | 1 | 283.681 | 15.072 | 87.928 | 0.146 | 0.138 |
| PD_all_dprime | 1 | 286.102 | 13.001 | 89.999 | 0.126 | 0.118 |
| Education | 1 | 299.715 | 0.414 | 102.586 | 0.004 | -0.006 |

```
+ FC_dprime


 Step 1 : AIC = 246.2373
 YN_dprime ~ FC_dprime
```

| Variable | DF | AIC | Sum Sq | RSS | R-Sq | Adj. R-Sq |
|---|---|---|---|---|---|---|
| ExecF | 1 | 230.905 | 9.417 | 51.927 | 0.496 | 0.486 |
| Age | 1 | 241.310 | 3.953 | 57.391 | 0.443 | 0.432 |
| PD_all_dprime | 1 | 244.691 | 2.057 | 59.287 | 0.424 | 0.413 |
| MoCA_Group | 1 | 244.813 | 1.987 | 59.357 | 0.424 | 0.412 |
| Education | 1 | 248.083 | 0.091 | 61.253 | 0.405 | 0.394 |

+ ExecF

Step 2 : AIC = 230.9053
YN_dprime ~ FC_dprime + ExecF

| Variable | DF | AIC | Sum Sq | RSS | R-Sq | Adj. R-Sq |
|---|---|---|---|---|---|---|
| Age | 1 | 228.245 | 2.276 | 49.651 | 0.518 | 0.503 |
| PD_all_dprime | 1 | 232.173 | 0.365 | 51.562 | 0.499 | 0.484 |
| MoCA_Group | 1 | 232.784 | 0.061 | 51.866 | 0.496 | 0.481 |
| Education | 1 | 232.885 | 0.010 | 51.917 | 0.496 | 0.481 |

+ Age

Step 3 : AIC = 228.2448
YN_dprime ~ FC_dprime + ExecF + Age

| Variable | DF | AIC | Sum Sq | RSS | R-Sq | Adj. R-Sq |
|---|---|---|---|---|---|---|
| PD_all_dprime | 1 | 229.835 | 0.195 | 49.456 | 0.520 | 0.500 |
| Education | 1 | 230.214 | 0.015 | 49.636 | 0.518 | 0.499 |
| MoCA_Group | 1 | 230.245 | 0.000 | 49.651 | 0.518 | 0.498 |

No more variables to be added.

Variables Entered:

+ FC_dprime
+ ExecF
+ Age

Final Model Output

### Model Summary

| | | | |
|---|---|---|---|
| R | 0.720 | RMSE | 0.705 |
| R-Squared | 0.518 | Coef. Var | 19764842892618133504.000 |
| Adj. R-Squared | 0.503 | MSE | 0.497 |
| Pred R-Squared | 0.479 | MAE | 0.524 |

RMSE: Root Mean Square Error
MSE: Mean Square Error
MAE: Mean Absolute Error

### ANOVA

| | Sum of Squares | DF | Mean Square | F | Sig. |
|---|---|---|---|---|---|
| Regression | 53.349 | 3 | 17.783 | 35.816 | 0.0000 |
| Residual | 49.651 | 100 | 0.497 | | |
| Total | 103.000 | 103 | | | |

### Parameter Estimates

| model | Beta | Std. Error | Std. Beta | t | Sig | lower | upper |
|---|---|---|---|---|---|---|---|
| (Intercept) | 0.000 | 0.069 | | 0.000 | 1.000 | -0.137 | 0.137 |
| FC_dprime | 0.474 | 0.078 | 0.474 | 6.086 | 0.000 | 0.320 | 0.629 |
| ExecF | 0.295 | 0.075 | 0.295 | 3.948 | 0.000 | 0.147 | 0.443 |
| Age | -0.163 | 0.076 | -0.163 | -2.141 | 0.035 | -0.315 | -0.012 |

### Selection Summary

| Variable | AIC | Sum Sq | RSS | R-Sq | Adj. R-Sq |
|---|---|---|---|---|---|
| FC_dprime | 246.237 | 41.657 | 61.343 | 0.40443 | 0.39859 |
| ExecF | 230.905 | 51.073 | 51.927 | 0.49586 | 0.48587 |
| Age | 228.245 | 53.349 | 49.651 | 0.51795 | 0.50349 |

3. Model selection for Forced Choice performance

```
Forward Selection Method
------------------------

Candidate Terms:

1 . Age
2 . Education
3 . MoCA_Group
4 . PD_all_dprime
5 . ExecF

 Step 0: AIC = 300.9723
 FC_dprime ~ 1
```

| Variable | DF | AIC | Sum Sq | RSS | R-Sq | Adj. R-Sq |
|---|---|---|---|---|---|---|
| MoCA_Group | 1 | 284.282 | 16.958 | 87.042 | 0.163 | 0.155 |
| Age | 1 | 286.590 | 15.024 | 88.976 | 0.144 | 0.136 |
| PD_all_dprime | 1 | 289.145 | 12.832 | 91.168 | 0.123 | 0.115 |
| ExecF | 1 | 291.458 | 10.802 | 93.198 | 0.104 | 0.095 |
| Education | 1 | 301.621 | 1.330 | 102.670 | 0.013 | 0.003 |

```
+ MoCA_Group


 Step 1 : AIC = 284.282
 FC_dprime ~ MoCA_Group
```

| Variable | DF | AIC | Sum Sq | RSS | R-Sq | Adj. R-Sq |
|---|---|---|---|---|---|---|
| Age | 1 | 277.499 | 6.984 | 80.057 | 0.230 | 0.215 |
| PD_all_dprime | 1 | 280.213 | 4.888 | 82.154 | 0.210 | 0.195 |
| ExecF | 1 | 283.329 | 2.414 | 84.628 | 0.186 | 0.170 |
| Education | 1 | 285.656 | 0.518 | 86.524 | 0.168 | 0.152 |

+ Age

Step 2 : AIC = 277.4993
FC_dprime ~ MoCA_Group + Age

| Variable | DF | AIC | Sum Sq | RSS | R-Sq | Adj. R-Sq |
|---|---|---|---|---|---|---|
| PD_all_dprime | 1 | 275.088 | 3.294 | 76.764 | 0.262 | 0.240 |
| ExecF | 1 | 277.582 | 1.448 | 78.609 | 0.244 | 0.222 |
| Education | 1 | 278.894 | 0.460 | 79.597 | 0.235 | 0.212 |

+ PD_all_dprime

Step 3 : AIC = 275.088
FC_dprime ~ MoCA_Group + Age + PD_all_dprime

| Variable | DF | AIC | Sum Sq | RSS | R-Sq | Adj. R-Sq |
|---|---|---|---|---|---|---|
| ExecF | 1 | 276.126 | 0.700 | 76.063 | 0.269 | 0.239 |
| Education | 1 | 276.173 | 0.666 | 76.098 | 0.268 | 0.239 |

No more variables to be added.

Variables Entered:

+ MoCA_Group
+ Age
+ PD_all_dprime

Final Model Output

### Model Summary

| | | | |
|---|---|---|---|
| R | 0.512 | RMSE | 0.872 |
| R-Squared | 0.262 | Coef. Var | 1712972823324994816.000 |
| Adj. R-Squared | 0.240 | MSE | 0.760 |
| Pred R-Squared | 0.202 | MAE | 0.658 |

RMSE: Root Mean Square Error
MSE: Mean Square Error
MAE: Mean Absolute Error

### ANOVA

| | Sum of Squares | DF | Mean Square | F | Sig. |
|---|---|---|---|---|---|
| Regression | 27.236 | 3 | 9.079 | 11.945 | 0.0000 |
| Residual | 76.764 | 101 | 0.760 | | |
| Total | 104.000 | 104 | | | |

### Parameter Estimates

| model | Beta | Std. Error | Std. Beta | t | Sig | lower | upper |
|---|---|---|---|---|---|---|---|
| (Intercept) | -0.468 | 0.199 | | -2.357 | 0.020 | -0.862 | -0.074 |
| MoCA_GroupPass | 0.600 | 0.230 | 0.249 | 2.608 | 0.010 | 0.144 | 1.056 |
| Age | -0.245 | 0.092 | -0.245 | -2.663 | 0.009 | -0.427 | -0.062 |
| PD_all_dprime | 0.194 | 0.093 | 0.194 | 2.082 | 0.040 | 0.009 | 0.379 |

### Selection Summary

| Variable | AIC | Sum Sq | RSS | R-Sq | Adj. R-Sq |
|---|---|---|---|---|---|
| MoCA_Group | 284.282 | 16.958 | 87.042 | 0.16306 | 0.15493 |
| Age | 277.499 | 23.943 | 80.057 | 0.23022 | 0.21512 |
| PD_all_dprime | 275.088 | 27.236 | 76.764 | 0.26189 | 0.23996 |
